# Federated analysis in COINSTAC reveals functional network connectivity and spectral links to smoking and alcohol consumption in nearly 2,000 adolescent brains

**DOI:** 10.1101/2022.02.02.478847

**Authors:** Harshvardhan Gazula, Kelly Rootes-Murdy, Bharath Holla, Sunitha Basodi, Zuo Zhang, Eric Verner, Ross Kelly, Pratima Murthy, Amit Chakrabarti, Debasish Basu, Subodh Bhagyalakshmi Nanjayya, Rajkumar Lenin Singh, Roshan Lourembam Singh, Kartik Kalyanram, Kamakshi Kartik, Kumaran Kalyanaraman, Krishnaveni Ghattu, Rebecca Kuriyan, Sunita Simon Kurpad, Gareth J Barker, Rose Dawn Bharath, Sylvane Desrivieres, Meera Purushottam, Dimitri Papadopoulos Orfanos, Eesha Sharma, Matthew Hickman, Mireille Toledano, Nilakshi Vaidya, Tobias Banaschewski, Arun L.W. Bokde, Herta Flor, Antoine Grigis, Hugh Garavan, Penny Gowland, Andreas Heinz, Rüdiger Brühl, Jean-Luc Martinot, Marie-Laure Paillère Martinot, Eric Artiges, Frauke Nees, Tomáš Paus, Luise Poustka, Juliane H. Fröhner, Lauren Robinson, Michael N. Smolka, Henrik Walter, Jeanne Winterer, Robert Whelan, IMAGEN Consortium, Jessica A. Turner, Anand D. Sarwate, Sergey M. Plis, Vivek Benegal, Gunter Schumann, Vince D. Calhoun

## Abstract

With the growth of decentralized/federated analysis approaches in neuroimaging, the opportunities to study brain disorders using data from multiple sites has grown multi-fold. One such initiative is the Neuromark, a fully automated spatially constrained independent component analysis (ICA) that is used to link brain network abnormalities among different datasets, studies, and disorders while leveraging subject-specific networks. In this study, we implement the neuromark pipeline in COINSTAC, an open-source neuroimaging framework for collaborative/decentralized analysis. Decentralized analysis of nearly 2000 resting-state functional magnetic resonance imaging datasets collected at different sites across two cohorts and co-located in different countries was performed to study the resting brain functional network connectivity changes in adolescents who smoke and consume alcohol. Results showed hypoconnectivity across the majority of networks including sensory, default mode, and subcortical domains, more for alcohol than smoking, and decreased low frequency power. These findings suggest that global reduced synchronization is associated with both tobacco and alcohol use. This work demonstrates the utility and incentives associated with large-scale decentralized collaborations spanning multiple sites.

## 1 Introduction

In the past decade, the field of neuroimaging has seen a rapid growth in data pooling/sharing initiatives (Eickhoff et al., 2016). This, in turn, has resulted in a major transformation in our understanding of brain disorders as well as accelerating studies to identify brain-based markers for potential use in clinical settings (Wilcox et al., 2020). However, open data sharing can be limited by policy or proprietary restrictions or data re-identification concerns (White et al., 2020) (Sweeney, 2002)(Sweeney, 2002, (Shringarpure and Bustamante, 2015). Such difficulties are further compounded in the case of sensitive patient data that cannot be easily shared or accessed. Finally, yet another challenge is pooling the data centrally can require significant computational resources which may not be available at a given site.

The aforementioned barriers can be offset by embracing federated analysis of neuroimaging data. Federated analysis does not require moving the data from the original sites but can generate results similar to pooled data. It can also be implemented with varying degrees of privacy, ranging from sharing of derived results only to full encryption. Towards this end, Plis et al. (2016) introduced a federated analysis framework and platform called the COllaborative Informatics and Neuroimaging Suite Toolkit for Anonymous Computation (COINSTAC; http://coinstac.trendscenter.org). There is no sharing of individual-level data in COINSTAC and thus the privacy of individual datasets is preserved while still allowing analyses to be performed. Multiple neuroimaging algorithms have been federated and can be run in COINSTAC. Examples of implemented algorithms include decentralized voxel-based morphometry (Gazula et al., 2018), decentralized t-distributed stochastic neighbor embedding (Saha et al., 2017, 2021), decentralized dynamic functional network connectivity (Baker et al., 2020), and decentralized support vector machine with differential privacy (Sarwate et al., 2014).

For the purpose of this study, we focus on decentralized variants of independent component analysis. Independent component analysis (ICA) is a versatile and powerful data-driven approach that can be used to analyze group fMRI data (Calhoun et al., 2009) to study the spatio-temporal structure of the fMRI signal. One of the earliest decentralized versions of ICA developed, for use with functional magnetic resonance imaging (fMRI) data, was the decentralized temporal ICA algorithm (Baker et al., 2015). More recently, (Baker et al., 2020) presented a decentralized group ICA algorithm to enable decentralized dynamic functional network connectivity analysis. Group ICA (Calhoun et al., 2009) enables group-level inferences while allowing for cross-subject variability. However, implementing group ICA within a decentralized context presents some challenges due to its fully data-driven output. For example, results typically require labelling of the output components, and while there are options for automated labelling (Salman et al., 2021) within the GIFT software (http://trendscenter.org/software/gift) the use of a data-driven approach involves considerable user interactions which can be inefficient in a federated environment. In addition, separate group ICA results may be challenging to compare with one another, hindering replication and cross-study comparison. To address this issue, Du et al. (2020) proposed the NeuroMark pipeline, an *a priori* driven and fully automated ICA approach informed by reliable network templates to achieve linked analyses among different datasets, studies, and disorders.

The decentralized Neuromark approach will be the subject of discussion in this work. Our focus is on the impact of substance use on adolescent brain functional network connectivity. Recently, Gazula et al. (2021) performed a large-N decentralized voxel-based morphometry analysis of structural magnetic resonance imaging data across two cohorts from 14 different sites to understand the structural changes in the brain as linked to age, body mass index and smoking. However, there has not yet been a large-N analysis of functional MRI data with ICA within a decentralized framework. In this work, we extend the aforementioned structural MRI results by presenting an analysis of functional network connectivity (FNC) and spectral analysis derived from independent component analysis on resting fMRI data. Our contributions in this paper can be summarized as follows.

1. Decentralized ICA on large datasets across multiple sites, IMAGEN from the Europe and cVEDA from India, in the COINSTAC framework and sharing observations.

2. Evaluating the association of smoking and alcohol use on functional network connectivity in the adolescent brain.

3. Decentralized MANCOVAN and univariate testing of ICA output.

The outline of the current paper is as follows: In Section 2, we discuss the Decentralized NeuroMark pipeline. In Section 2.3, we describe the IMAGEN and cVEDA data used in this study. In Sections 3 and 4, we present the results and discuss our experience implementing the Neuromark pipeline in COINSTAC. We conclude the paper in Section 5.

## 2 Methods & Data

As mentioned earlier, the goal of group ICA is to enable group level inferences by allowing for subject-level variability. However, due to the data-driven nature of ICA, these inferences can turn out to be inconsistent when compared across different studies containing data from similar disorders. It is possible to systematically integrate these findings across studies and disorders by interlinking the estimation of subject-level spatial maps using a common set of spatial priors and Neuromark (Du et al., 2020) precisely achieves this objective. The goal of Neuromark is to fully automate the estimation and labeling of individual subject connectivity features using spatial network priors derived from independent large samples. Another advantage of Neuromark is that the ICA estimates are computed on individual subjects, which completely avoids data leakage, facilitating cross-validation and independent analysis. The estimation of subject specific networks is performed by leveraging spatially-constrained ICA (Lin et al., 2010; Du and Fan, 2013).

The Neuromark pipeline leverages component priors derived from replicable spatial maps estimated from large independent datasets. For this purpose, resting-state fMRI datasets from the Human Connectome Project (HCP, http://www.humanconnectomeproject.org/data/) and the Genomics Superstruct Project (Holmes et al., 2015) were utilized. Group ICA was applied separately to the two datasets, followed by a greedy spatial correlation analysis (coupled with expert inspection) to identify and label intrinsic connectivity networks (ICNs) that showed high replicability across the two separate group ICA analyses (Du et al., 2020).

For the Neuromark pipeline, these ICNs are used as network templates (aka priors) to estimate subject-specific functional networks and their associated time-courses (TCs) via an adaptive spatially constrained ICA method. Here we used multi-objective optimization ICA with reference where one objective is to optimize the independence of networks (for the subject), while the other is to optimize the similarity between one subject-specific network and its related network template. Once this is accomplished, various network features can be extracted from both static and dynamic perspectives. One such example of a measure is the static FNC (sFNC) which is computed as the Pearson correlations between TCs of ICNs to yield an sFNC matrix reflecting the interaction between any two networks. While the spatial map of each ICN reflects intra-connectivity within brain functional network, sFNC matrix represents inter-connectivity strengths between different ICNs. The obtained network features and time courses can be further analysed statistically via decentralized MANCOVA for making inferences.

### 2.1 Decentralized ICA stats

Once the network features are derived from Neuromark, the researcher/user can perform any statistical analysis in a decentralized fashion. This helps in speeding up the analysis enabling multiple research groups to participate in larger studies, preserving their data privacy without sharing the actual data. Fig. 1 shows how the Neuromark pipeline is performed in a centralized way. There are two ways in which the decentralized statistics can be performed. Here we implement the MANCOVAN toolbox in GIFT (https://trendscenter.org/software/gift/), which provides both MANCOVAN omnibus tested as well as univariate regression tests. In this work, we utilize the univariate regression functionality, though the MANCOVAN toolbox (MTB) is fully implemented. The first approach, *d*MANCOVAN (decentralized MANCOVA), involves pooling all the time courses within a private aggregator and running the MTB on these pooled data following which the results are shared with every local site (Algorithm 1 and Figure 2). The second and more secure approach dMANCOVA*pa* (decentralized MANCOVA with private aggregator) involves performing an MTB analysis at each local site separately on its corresponding timecourses and covariate data. From the local MTB results generated, relevant stats information such as residual errors, contrast mean images, and other metadata (such as number of subjects) are extracted and sent to the master site. At the master site, these stats and metadata from all the local sites are combined to compute global sum of squares residual errors and average beta weights. From these, t-values and p-values are computed and these results are sent to all the local sites. Alg. 2 and Figure 3 show the algorithm and pipeline corresponding to the improved decentralized MTB approach respectively.

**Figure 1:**
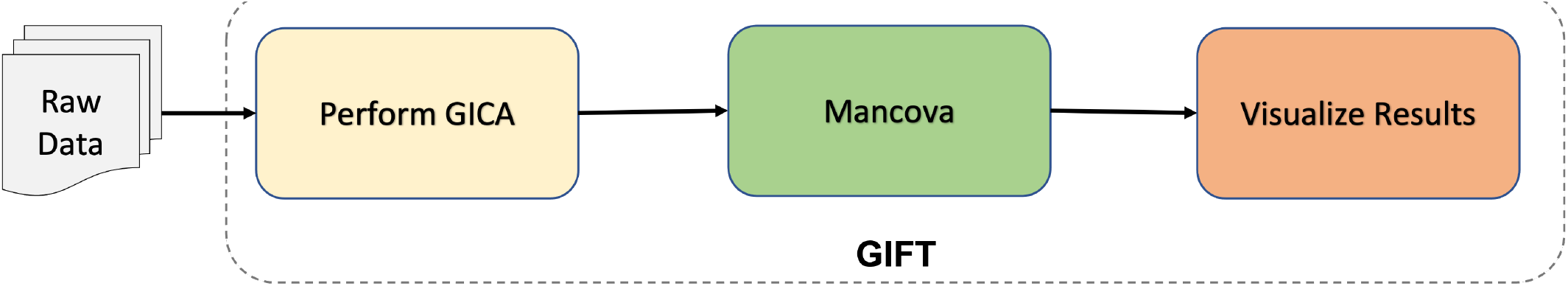
Pipeline of GIFT framework computed in a centralized way

**Figure 2:**
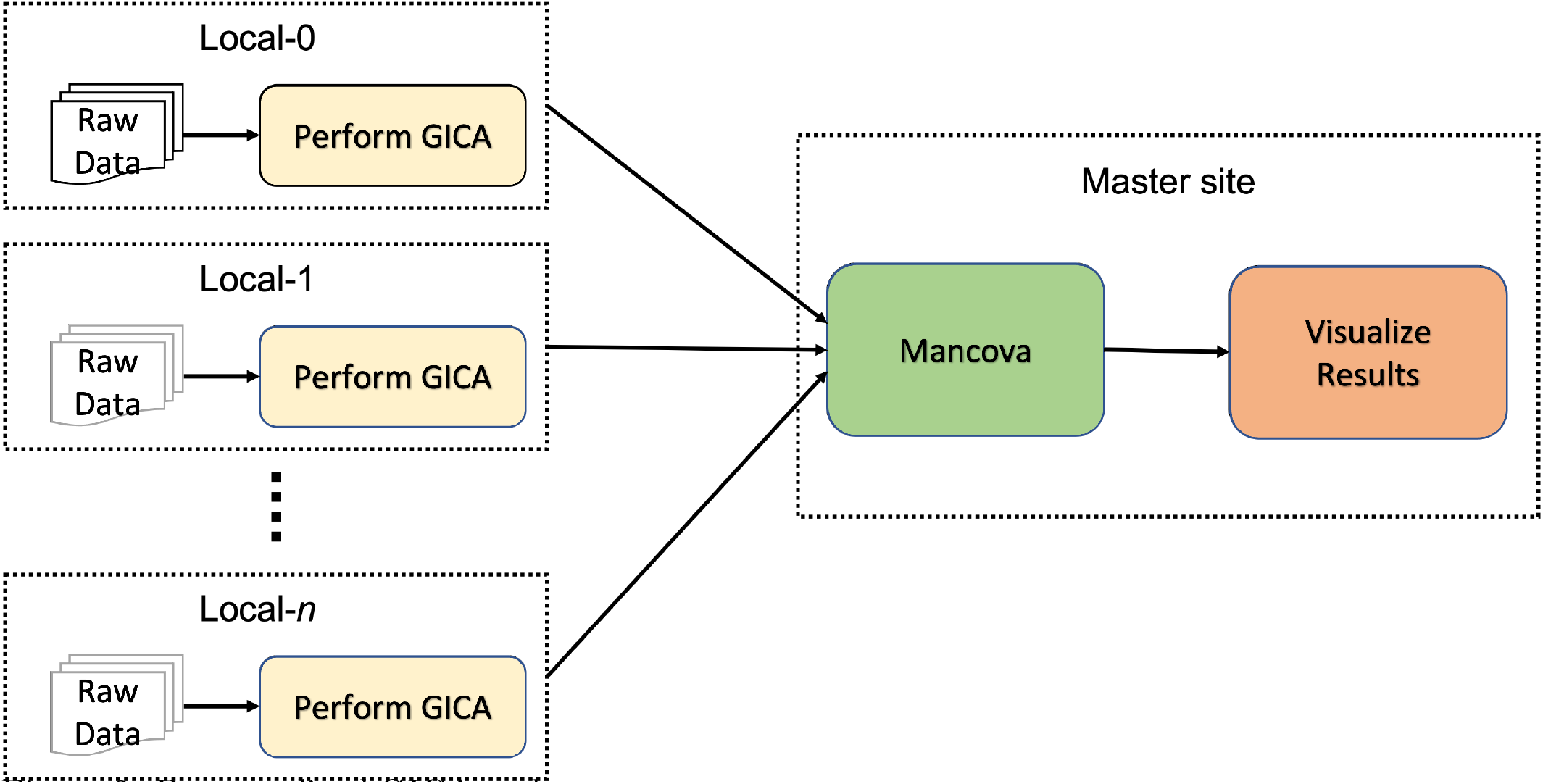
Decentralized GICA and mancova implemented in COINSTAC using GIFT toolbox. This approach involves pooling all the time courses within a private aggregator and running the MTB on this pooled data following which the results are shared with every local site.

**Figure 3:**
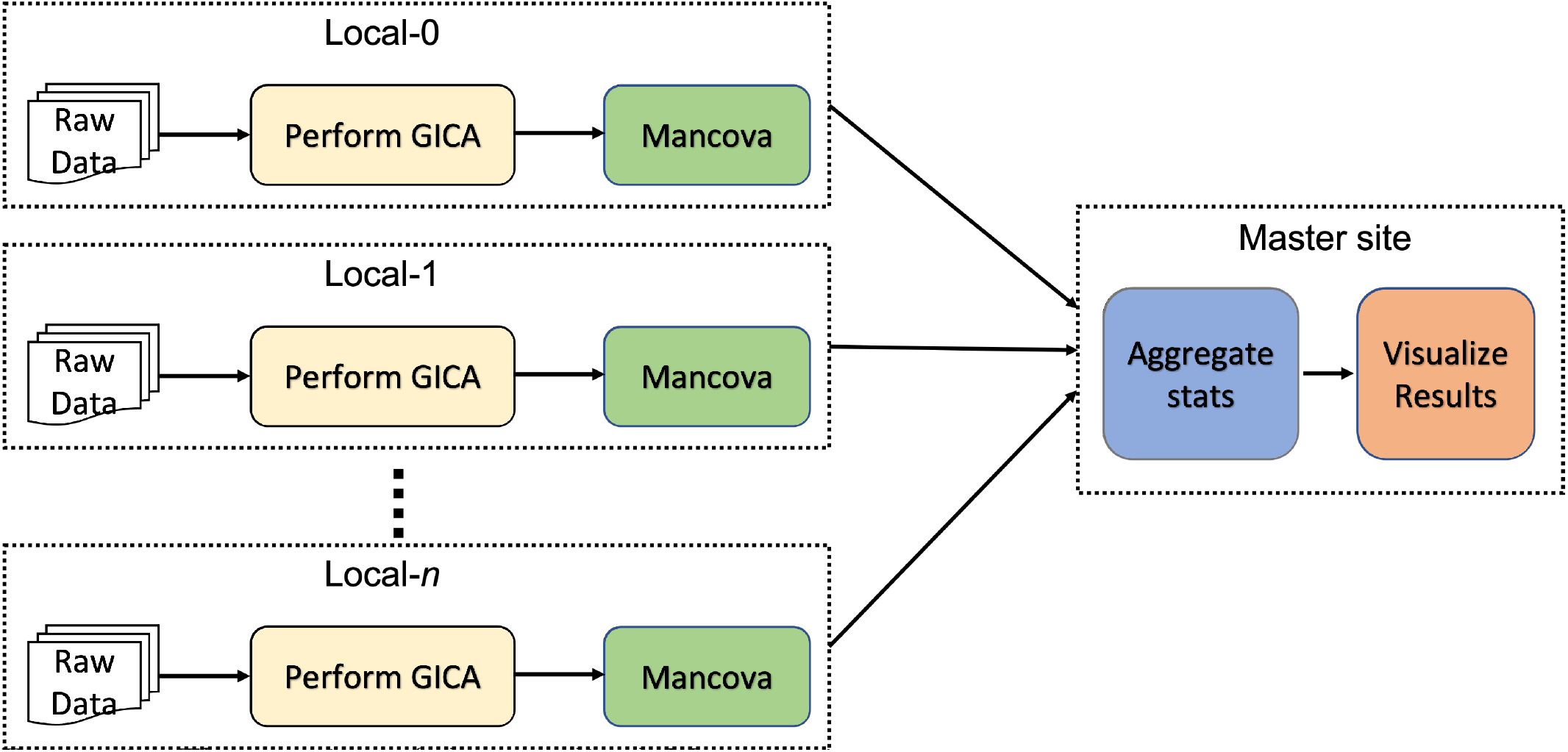
This version of decentralized Mancova involves performing an MTB analysis at each local site separately on its corresponding timecourses and covariate data. From the local MTB results generated, relevant stats information such as residual errors, contrast mean images, and other metadata (such as number of subjects) are extracted and sent to the master site. At the master site, these stats and metadata from all the local sites are combined to compute global sum of squares residual errors and average beta weights. From these, t-values and p-values are computed and these results are sent to all the local sites.

**Figure 4:**
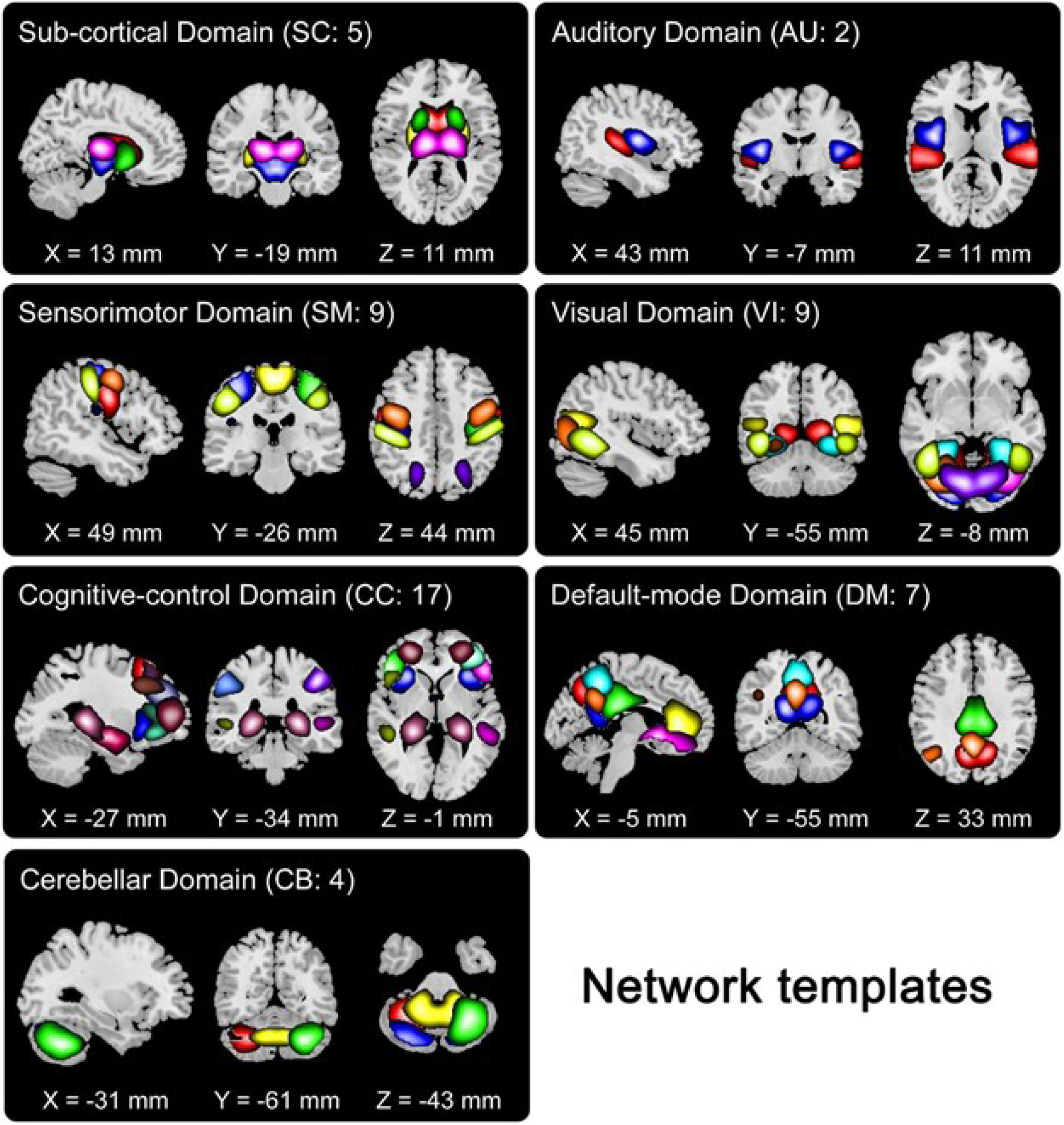
Domains identified in Du *et al*. Briefly, these seven identified network templates were divided based on anatomical and functional properties. In each subfigure, one color in the composite maps corresponds to an intrinsic connectivity network (ICN).

**Figure 5:**
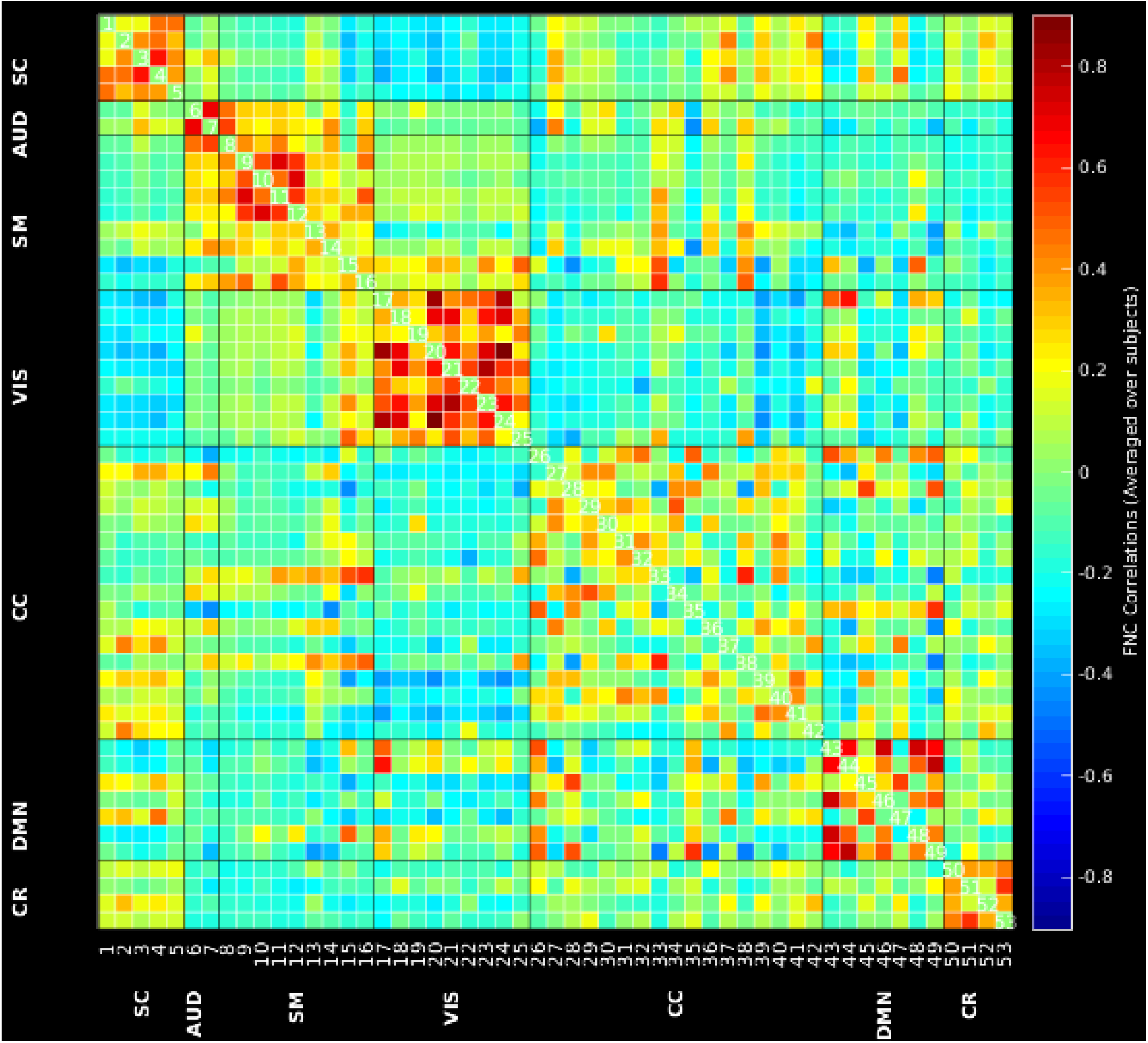
Correlation map showing average correlation, across participants, for each domain. Cool colors represent anticorrelation and warm colors represent positive correlation. In general, there was high positive (r ¿ .5) correlation within the visual, sensorimotor, and default mode network domains across all participants. Of note. SC: sub-cortical domain, AUD: auditory domain, SM: sensorimotor domain, VIS: visual domain, CC: cognitive-control domain, DMN: default-mode domain, CR: cerebellar domain.

#### Algorithm 1 Decentralized GICA and MANCOVA

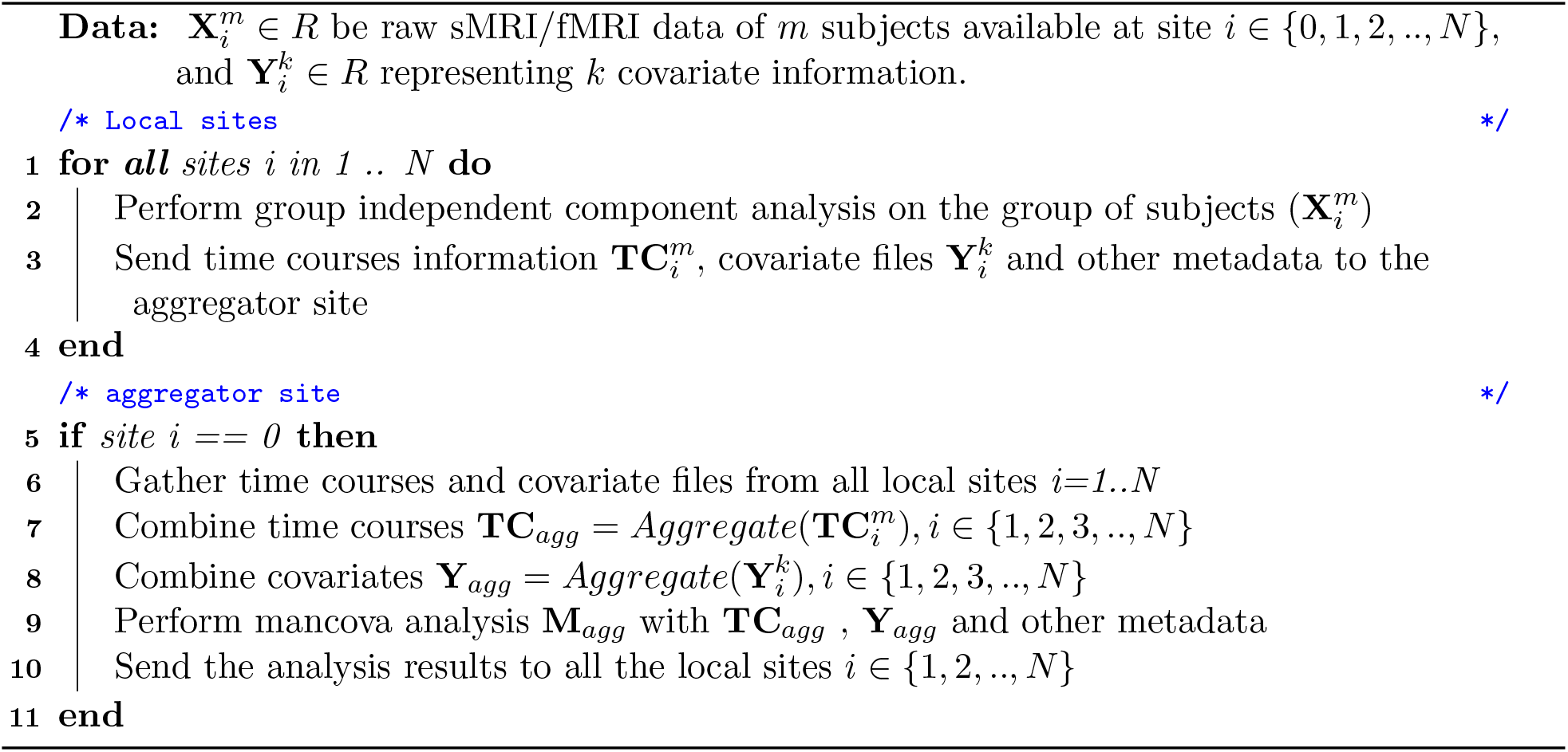

## 2.2 COINSTAC

For this project, the entire analysis, neuromark and the decentralized statistical analysis, was performed in COINSTAC. In brief, COINSTAC (Plis et al., 2016; Ming et al., 2017; Gazula et al., 2020) is a fully open-source, federated-learning platform geared towards neuroimaging research enabling researchers to perform decentralized analysis of neuroimaging data in a secure and privacy-preserving environment via a message-passing infrastructure. More specific instructions on how to install the application and run an example analysis can be found online at https://github.com/trendscenter/coinstac-instructions.

### Algorithm 2 Decentralized GICA and MANCOVA with Private Aggregator

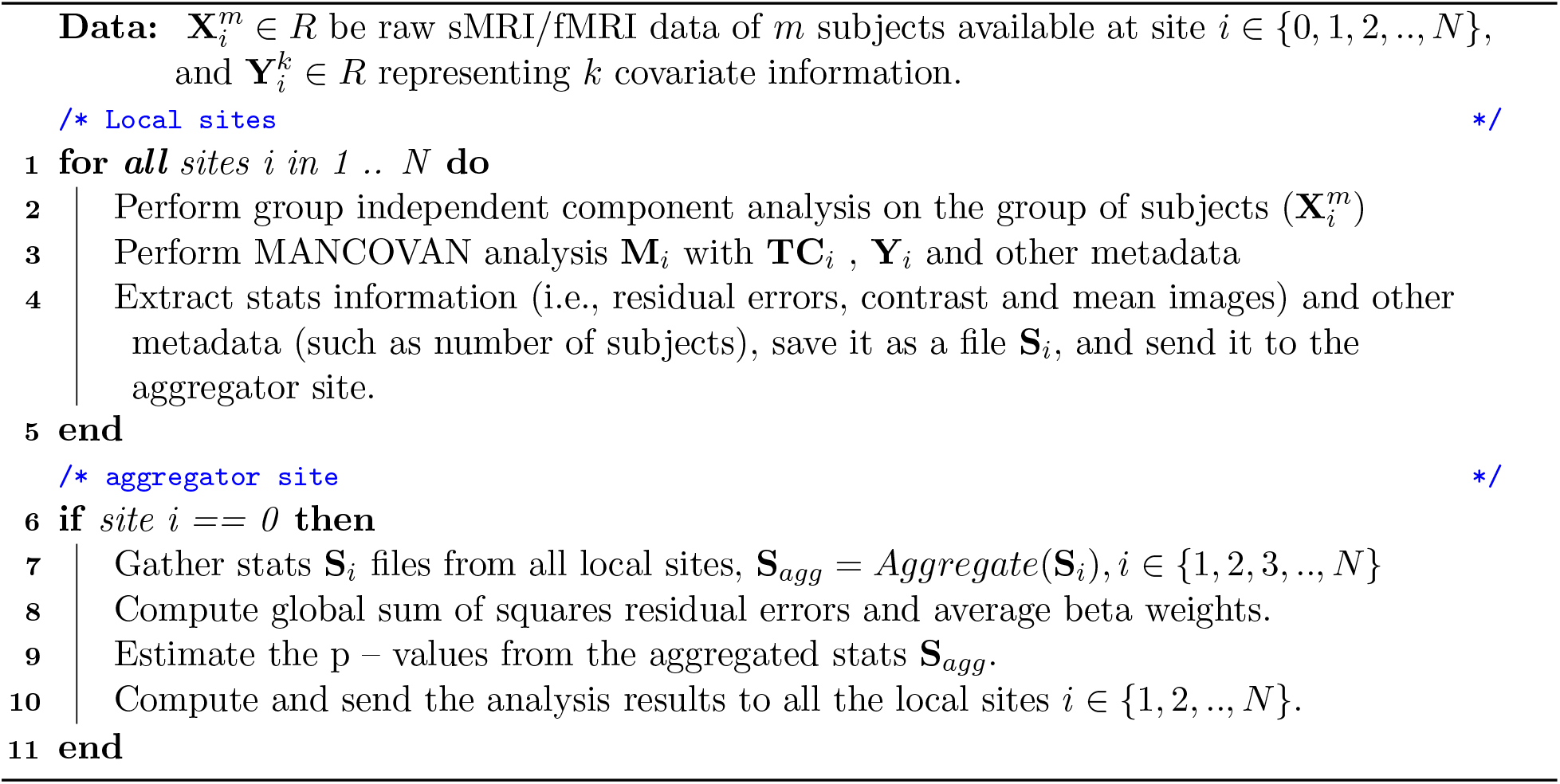

### 2.3 Data

#### 2.3.1 Cohort

The data from two large multi-centric adolescent cohorts in India and Europe were used for the present analysis. The Consortium on Vulnerability to Externalising Disorders and Addiction (cVEDA) is an accelerated longitudinal cohort in India (Sharma et al., 2020) that covers an age span of 5-24 years. Participants who provided brain imaging data were recruited at six study sites (see Zhang et al. (2020) for cohort profile and Sharma et al. (2020) for study protocol). The IMAGEN project is a longitudinal study of adolescent brain development and mental health in Europe (Schumann et al., 2010). Participants were recruited from eight study sites in England, Ireland, France, and Germany at the age of 14, with follow-up assessments at ages 16, 19, and 22. Each study site obtained ethical approval from the local research ethics committee. For both studies written consent (or verbal) assent were acquired from all the participants and parents prior to participation.

In this research, we used resting functional neuroimaging, and smoking and drinking assessments from the baseline cVEDA cohort (n=1140) and the second follow-up wave in IMAGEN (n=839), acquired at age 19. See Table 1 for the cohort description.

**Table 1:** Sample Characteristics

| **Variable** | **cVEDA (N=1140)** | **IMAGEN (N=839)** | **Total (N=1979)** |
| --- | --- | --- | --- |
| Age [y, Mean (SD)] | 15.677 (4.346) | 18.998 (0.718) | 17.085 (3.714) |
| Sex [Male, n (%)] | 638 (56.0%) | 389 (46.4%) | 1027 (51.9%) |
| BMI [kg/m<sup>2</sup>, Mean (SD)] | 19.237 (4.509) | 22.836 (3.946) | 20.763 (4.634) |
| Smoking Risk, n (%): 0 | 1073 (94.1%) | 242 (28.8%) | 1315 (66.4%) |
| 1 | 67 (5.9%) | 597 (71.2%) | 664 (33.6%) |
| Alcohol Risk, n (%): 0 | 1088 (95.4%) | 224 (26.7%) | 1312 (66.3%) |
| 1 | 43 (3.8%) | 516 (61.5%) | 559 (28.2%) |
| 2 | 9 (0.8%) | 99 (11.8%) | 108 (5.5%) |

#### 2.3.2 MRI dataset acquisition and preprocessing

Although COINSTAC also implements fMRI preprocessing, for this study data was previously preprocessed by each site separately outside of COINSTAC, and we focused on using COINSTAC to implement a group ICA + stats analysis. The preprocessing steps are briefly described below. For a complete description of the preprocessing of the IMAGEN dataset see Schumann et al. (2010) and for the cVEDA dataset see (Sharma et al., 2020; Zhang et al., 2020).

For IMAGEN, data preprocessing was performed using the statistical parametric mapping (SPM) tool (http://www.fil.ion.ucl.ac.uk/spm/) toolbox. fMRI BOLD images were first realigned, corrected for head motion and slice timing (to account for timing difference in slice acquisition) and co-registered to T1w (MPRAGE) images. The fMRI data were normalized to the EPI template and resampled to 3 mm^3^ isotropic voxels. Images were spatially smoothed using a Gaussian kernel with a 6 mm full width at half maximum. Subjects with head motion < 0.25 mm (frame-wise displacement) and with functional data providing near full brain successful normalization (by comparing the individual mask with the group mask) were selected for further analysis.

For cVEDA, the parameters and acquisition protocols were similar to those described above in IMAGEN. More detailed information can be found at https://cveda-project.org/standardoperating-procedures/. The functional data processing for cVEDA in brief included the following steps. Motion correction by applying a rigid body registration of each volume to the middle volume (FSL MCFLIRT). Slice-timing correction to account for timing difference in slice acquisition. Non-brain tissue removal (FSL BET) and co-registration to high-resolution T1 image (FSL FLIRT using the BBR algorithm). Motion correcting transformations, BOLD-to-T1w transformation and T1w-to-template (MNI) warp were concatenated and applied in a single step using Advanced Normalization Toolbox (ANTs v2.1.0) and Lanczos interpolation. An ICA-based Automatic Removal Of Motion Artifacts (AROMA) was used to generate non-aggressively denoised data. And finally these data was resampled to 3mm isotropic and smoothed using a 6mm non-linear filter using FSL SUSAN. Only those subjects who had head motion FD of <0.25mm were retained.

#### 2.3.3 Measure of Tobacco and Alcohol use

For IMAGEN, measures of tobacco smoking were obtained from self-reports in the Fagerstrom test for nicotine dependence (FTND) questionnaire (Heatherton et al., 1991). The answer to the question regarding lifetime smoking experience was used to create a binary variable, indicating whether the participant had ever smoked (labeled as 1) or not (labeled as 0). For alcohol risk, the Alcohol Use Disorders Identification Test–Consumption (AUDIT-C) score < 3 for females or < 4 for males were labeled as 0 (low risk), score of 3–7 for females or 4–7 for males were labeled 1 (moderate risk) and score *≥* 8 was labelled as 2 (high risk).

For cVEDA, measures of tobacco smoking and alcohol use were obtained from the World Health Organization’s Alcohol, Smoking and Substance Involvement Screening Test (ASSIST) questionnaire, which provides a specific substance involvement score indicating the risk levels. For smoking risk, a tobacco involvement score of > 3 was labeled as 1 (present) and a score of *≤* 3 was labeled as 0 (absent). For alcohol risk, an involvement score of *≤* 3 was labeled 0 (low risk), a score 4–26 was labeled 1 (moderate risk) and *≥* 27 was labeled as 2 (high risk).

Other covariates from both the cohorts were mean-centered indices of head motion during resting state functional MRI (measured as framewise displacement in mm), body mass (BMI) (weight in kilograms divided by the squared height in meters), age in years, and dummy-codes for site and sex. (refer to Table 1 for sample characteristics).

## 3 Results

We provide a summary of the results here and then discuss them further in the following section. Please note that all the maps have been thresholded with a false discovery rate (FDR) correction of 0.05 (Benjamini and Hochberg, 1995).

### 3.1 FNC for Alcohol

Lifetime risk for alcohol showed more mean functional connectivity strengths than lifetime risk for tobacco. Across the seven domains (default mode network (DMN), sub-cortical (SC), auditory (AUC), sensorimotor (SM), visual (VIS), cognitive-control (CC), & cerebellar (CR)), there were roughly 27 positive pairs and 48 negative pairs, the SC domain was most positively associated with the other domains (8 positive pairs) and the SM domain and the CC domains were the most negatively associated with the other domains (20 and 21 pairs, respectively). Increased functional connectivity was observed between the DMN and cognitive control. See Figure 6 for more details.

**Figure 6:**
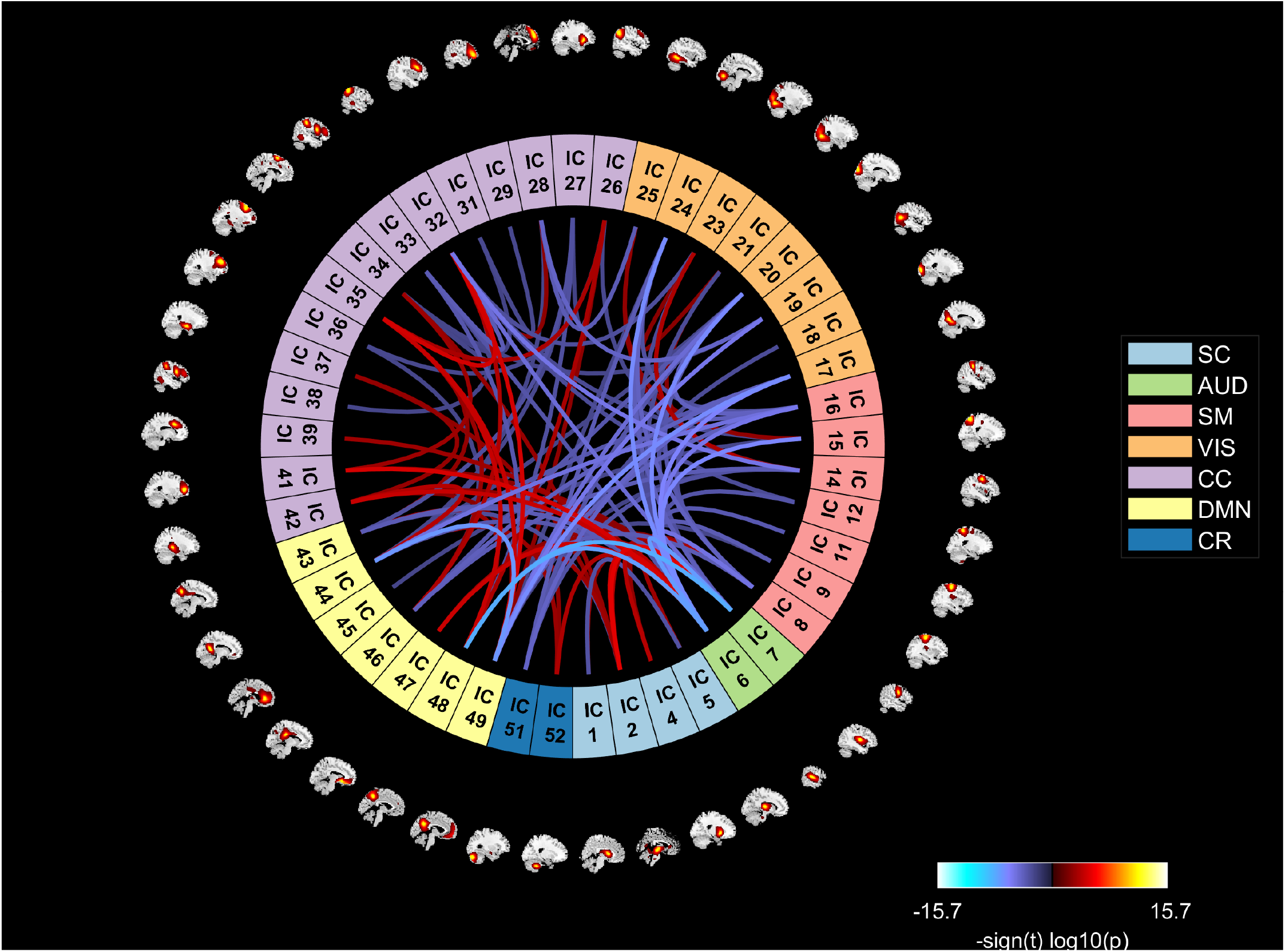
Connectogram showing group mean connectivity across domains for alcohol users. Cool colors represent reduced connectivity and warm colors represent increased connectivity. In alcohol users, there was increased connectivity mostly linked to the DMN domains and to a lesser extent, the cognitive control and sub-cortical domains. Of note. SC: sub-cortical domain, AUD: auditory domain, SM: sensorimotor domain, VIS: visual domain, CC: cognitive-control domain, DMN: default-mode domain, CR: cerebellar domain.

### 3.2 FNC for Tobacco

In addition to less overall connectivity strength, lifetime risk for tobacco showed almost exclusive negative connectivity pairs across the seven domains. To a lesser extent than alcohol, the DMN domain had increased functional connectivity with the cognitive control domain, a total of four positive connectivity pairs. The DMN also had one positive connectivity pair with the cerebellar domain and the visual domain had one positive connectivity pair with the sensorimotor domain. All of the other 27 connectivity pairs were negatively associated. See Figure 7 for more details.

**Figure 7:**
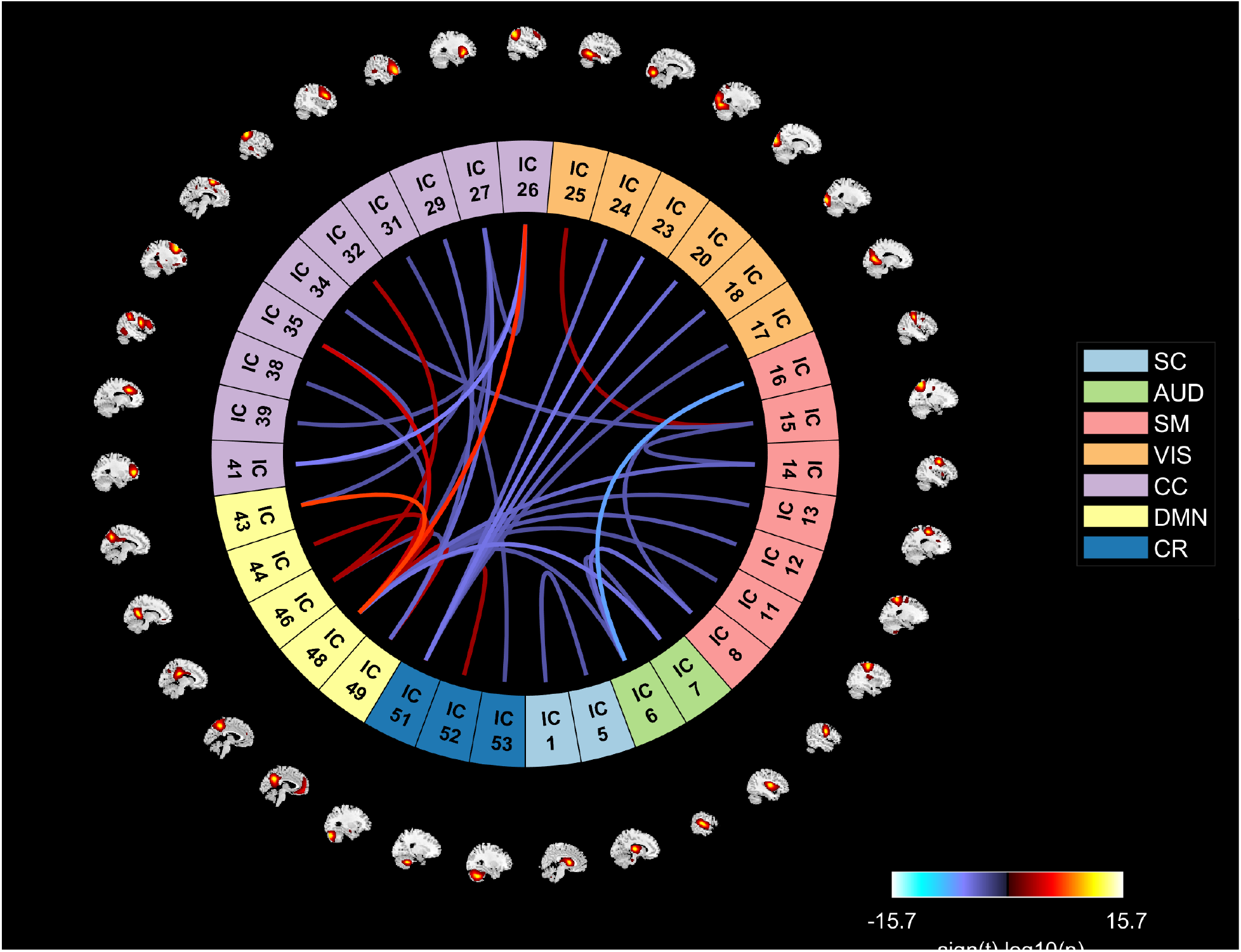
Connectogram showing group mean connectivity across domains for tobacco users. Cool colors represent reduced connectivity and warm colors represent increased connectivity. In tobacco users, there was decreased connectivity linked to sensorimotor, auditory, and visual domains. Of note. SC: sub-cortical domain, AUD: auditory domain, SM: sensorimotor domain, VIS: visual domain, CC: cognitive-control domain, DMN: default-mode domain, CR: cerebellar domain.

For both lifetime risk of alcohol and tobacco, there was a predominant decrease in a number of the connections, suggesting a common reduction of connection effect for both substances. Taken together, there was a decreased functional network connectivity observed in the sensorimotor, auditory, and visual networks in both alcohol and tobacco use.

The univariate results showed reduced power in the high frequency region for lifetime alcohol and tobacco use (both above 0.15 Hz) with larger changes for alcohol than tobacco. Alcohol use also showed increased power in the mid/BOLD frequency range, see Figures 8 and 9 for more details. As a whole, alcohol use indicated two sets of signal frequencies, reduced power (roughly 0.2 Hz) at high frequency and high power (between 0.05 - 0.1 Hz) at low frequency.

**Figure 8:**
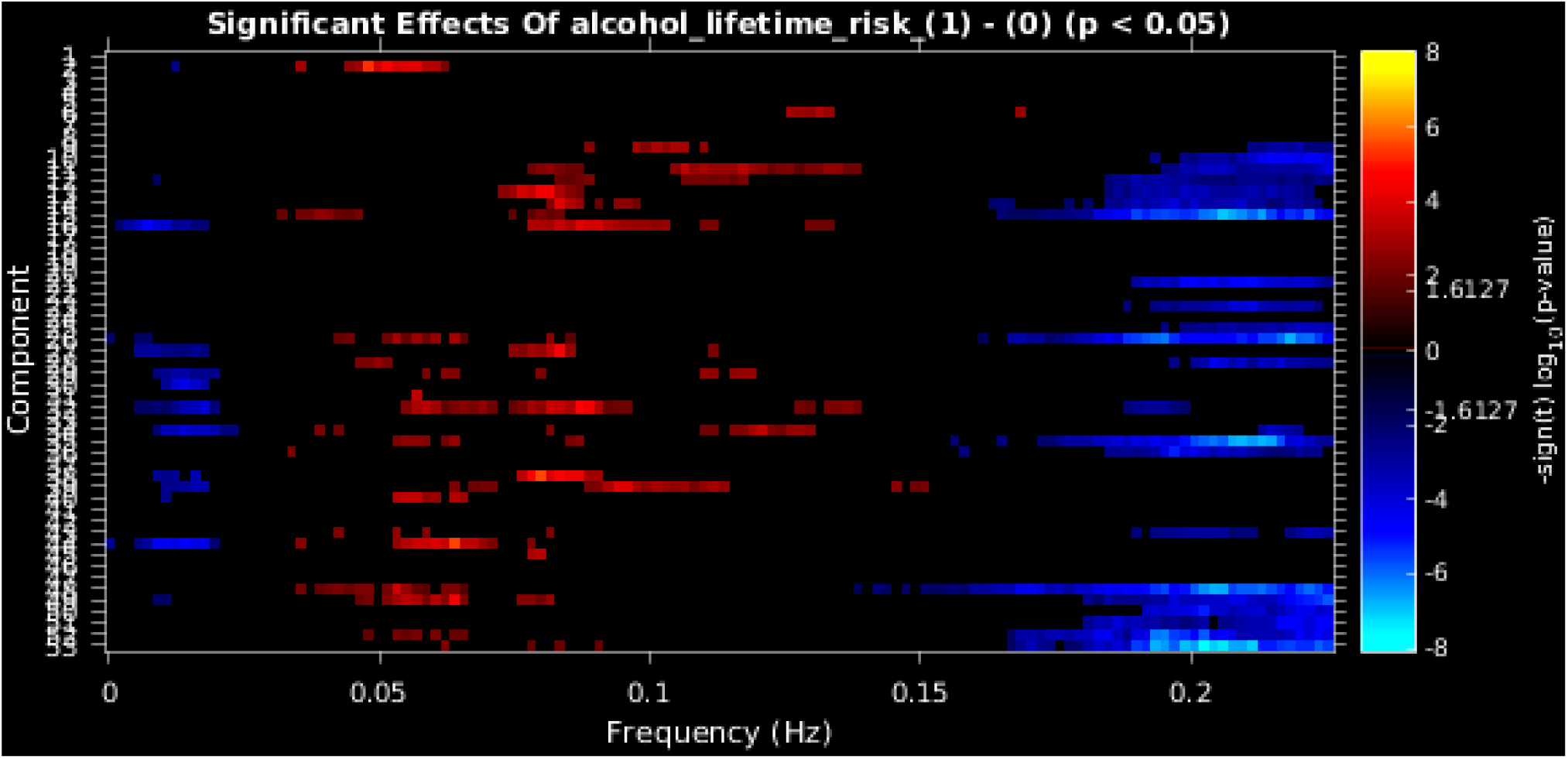
Power spectra of each component for alcohol risk (condition 1) vs no risk (condition 0) flagged red or blue based on the direction of effect for every value where *p* ¡ 0.05. There is a general increase in the mid/BOLD frequency range (between 0.05 - 0.1 Hz) and reduced high frequency power above 0.15 Hz.

**Figure 9:**
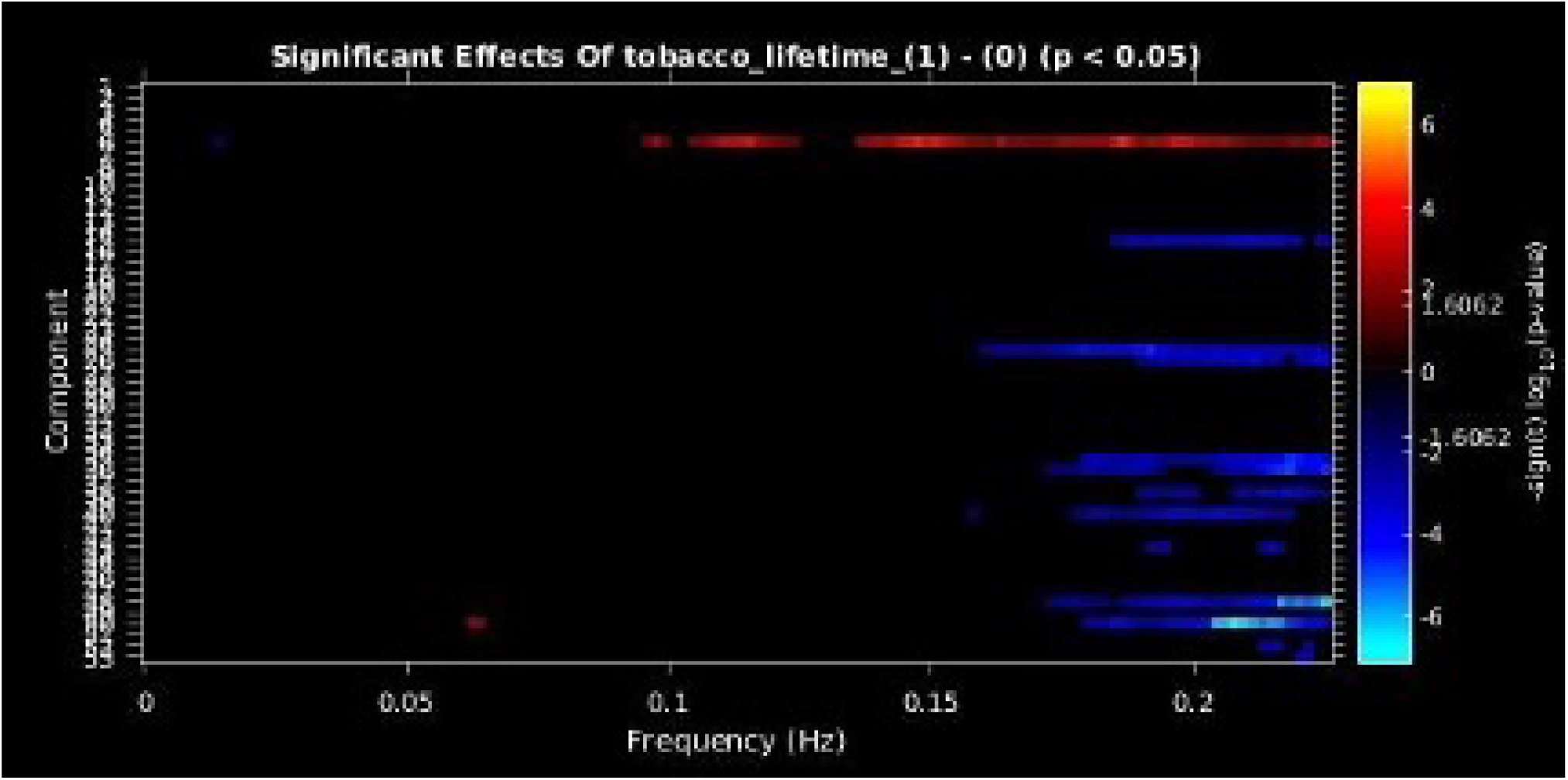
Power spectra of each component for effect of tobacco risk (condition 1) vs no risk (condition 0) flagged red or blue based on the direction of effect for every value where *p* ¡ 0.05. There was reduced BOLD high frequency power (above 0.15 Hz) across all components.

#### 3.2.1 Decentralized MANCOVA

We test our approach using the subset of the fMRI dataset used in Allen et al. (2011). Since, it is a large dataset, we select a subset of 50 subjects and run mancova algorithm as shown in Fig. 1. The same data is partitioned into two subsets to test our decentralized mancova analysis using algorithms Alg. 1 and Alg. 2. In our results, we compare FNC correlations and significant effects of gender outputs across different algorithms.

Fig. 10, 11, 12 show the FNC correlations, significant effects of gender and effect sizes per gender. These are the general studies performed using mancova analysis. From the figures, it can be seen that Alg. 1 has same output as the GIFT interface in all the figures. Alg. 2 has the same output for FNC correlations (Fig. 10(b), 10(c)) but varies slightly in terms of significant effects of gender (Fig. 11(b), 11(c) and Fig. 12(b), 12(c)). This is because the mancova output from all the local sites were used in computing mancova output in Alg. 2 rather than computing mancova at the remote site as in Alg. 1. This improvisation in Alg. 2 reduces running time by 50% as compared to Alg. 1.

**Figure 10:**
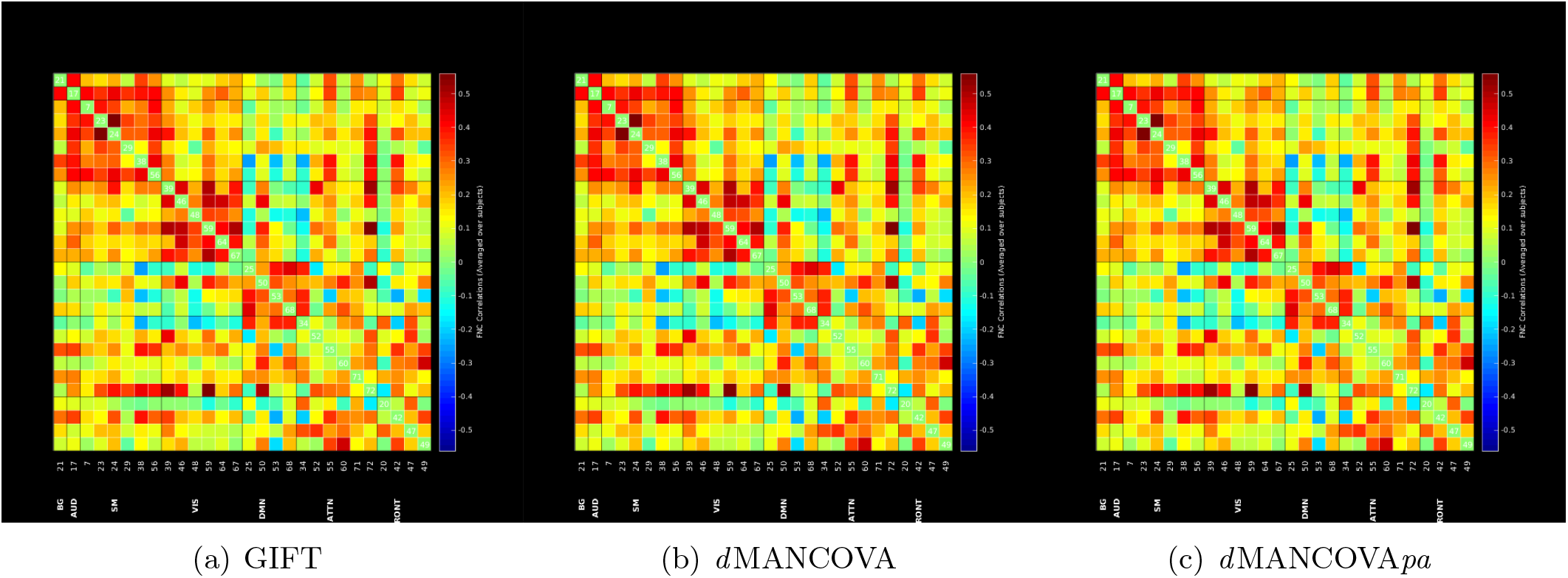
Functional Network Connectivity matrices generated using GIFT interface, *d* MANCOVA (Alg. 1) and *d* MANCOVA*pa* (Alg. 2)

**Figure 11:**
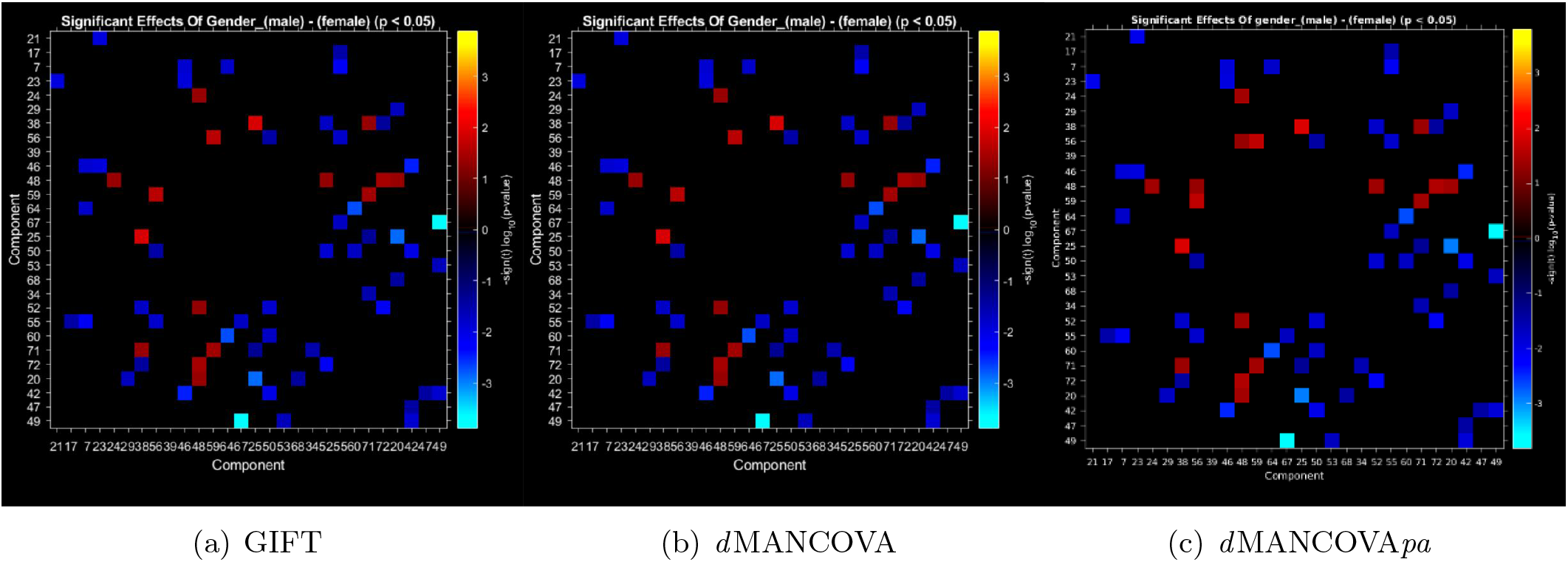
Significant effects of gender generated using GIFT interface, *d* MANCOVA (Alg. 1) and *d* MANCOVA*pa* (Alg. 2).

**Figure 12:**
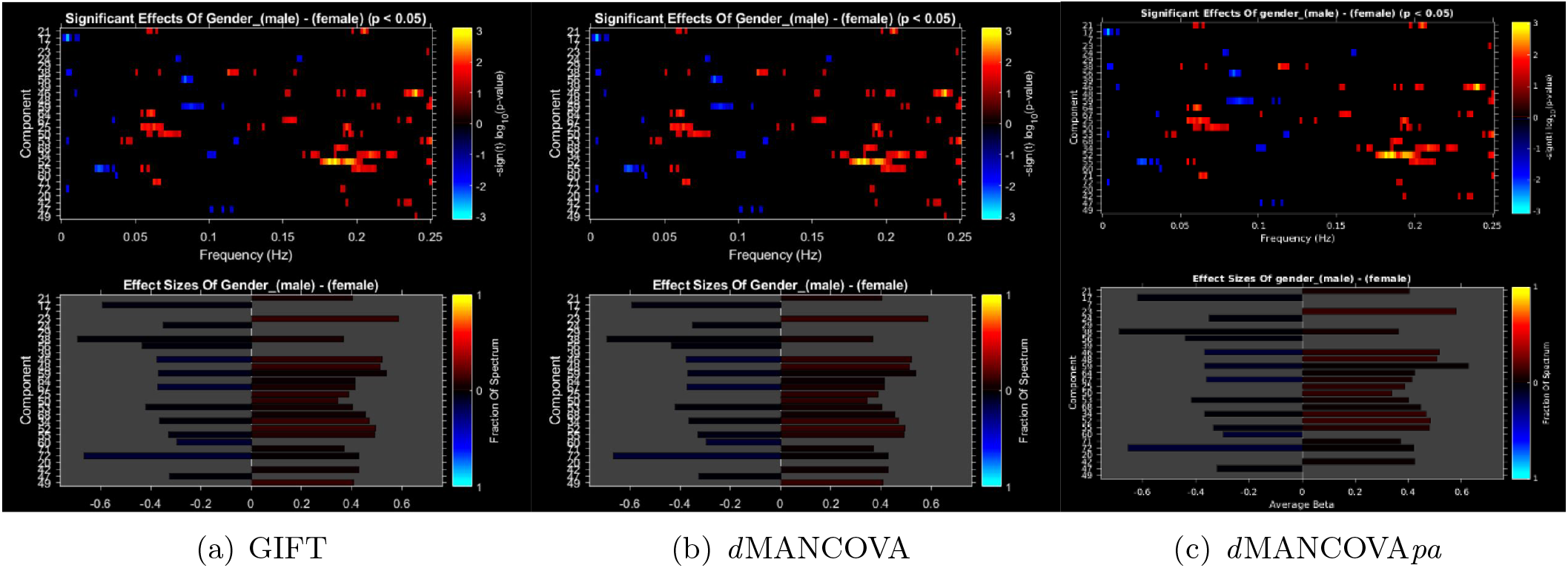
Significant effects and effect sizes of gender generated using GIFT interface, dMANCOVA (Alg. 1) and *d* MANCOVA*pa* (Alg. 2).

## 4 Discussion

The goal of this paper was two-fold: to demonstrate the feasibility of COINSTAC (Gazula et al., 2018) for performing decentralized analysis on large datasets present across multiple sites, compare the results of the decentralized ICA and Neuromark, and to use this experiment to evaluate the relationships of both smoking and tobacco use with functional connectivity changes. Using a large sample of tobacco and alcohol users, this study investigated resting state functional connectivity in seven brain networks (SC, AUD, SM, VIS, CC, DMN, & CR). Using previously-defined networks (Du et al., 2020) we identified a trend of general decreased connectivity across all networks in both tobacco and alcohol users.

Starting with tobacco users, there was decreased connectivity in most of the pairings but the most prominent result was hypoconnectivity between the auditory and sensorimotor networks. These findings are in support of the reduction of global cortical efficacy in smokers (Fedota and Stein, 2015). There was an exception with the DMN, that had increased within-network connectivity but also with the cognitive control and cerebellar domains. The increased connectivity between the DMN, which is associated with self-referential processes (Andrews-Hanna et al., 2010) and the cerebellum may be related to associative reward learning and cravings as the cerebellar domain has been previously associated with encoding of predicted internal states (Ebner and Pasalar, 2008; Popa and Ebner, 2019). Additionally, hyperconnectivity specifically within the DMN has been noted before (Vergara et al., 2017) and was hypothesized as an indication of the body being in a state of withdrawal. A similar effect was identified in cannabis users (Pujol et al., 2014). Therefore, we speculate that our results of increased connectivity may be related to states of cravings or withdrawal (e.g., time in the scanner prevented participants from smoking) and not a direct connectivity alteration relating to tobacco use.

In alcohol users, there was significantly more connectivity across all networks when compared to tobacco users. Generally, there was hyperconnectivity between the DMN, CC, and SC domains. One explanation for this increased connectivity may be compensation related to impaired functionality that results from prolonged alcohol use. In other words, the hyperconnectivity may be a sign of overactivation, or enhanced neural effort, that is needed to achieve the original (or baseline) cortical performance in areas that have been negatively affected by alcohol (Jansen et al., 2015; Zhu et al., 2017). As some of these hyper-connections (within DMN and DMN and CC) were seen with the tobacco users, another potential explanation is the previously stated hypothesis that the hyperconnectivity is representing a state of withdrawal and not a direct change from alcohol use. However, these results would benefit from future studies examining how much of this connectivity increase is related to the stages (and degree) of withdrawal or if this increase is still identified after substance consumption.

The remainder of the 75 identified pairs in alcohol use indicated hypoconnectivity. The negative pairs were across all seven domains, showing support for global cortical deficits in alcohol users. The strongest of those were within the DMN and between the DMN and AUD domains and between AUD and VIS domains. The visual region, specifically the middle occipital gyrus (Camchong et al., 2012), has previously been identified as a primary area of impact in alcohol users (Vergara et al., 2017). The hypoconnectivity between the DMN and VIS indicate a reduction in domains known for self-reference and information processing related to one’s own body, aspects that when not functioning properly could explain excessive consumption, among other decisions.

Finally, both tobacco and alcohol users had general hypoconnectivity within the DMN, SM, VIS, AUD, CC, and CR domains. Alcohol users had more connections than tobacco users, with the majority of those pairs also being negative. These findings suggest that global reduced synchronization is associated with both tobacco and alcohol use. Specific domains, namely the DMN, CC, and VIS domains may explain some of the mechanisms behind substance use and how addictions are formed.

The results of the decentralized MANCOVA with a private aggregrator show that it is possible to obtain results similar to results where the analysis is performed locally. In fact, it can also be seen that the results also qualitatively match the ones from a centralized analysis. This is a good indicator of how end-to-end large scale pipelines can be successfully implemented in COINSTAC without concerns about loss of significance.

In summary, we proposed an ICA-based framework to generalize and standardize the calculation of possible functional connectivity features that leverages the benefits of a data-driven approach and also provides comparability across multiple analyses. In the current study, the experimental conditions for data collection and preprocessing is specific to each site thus enabling a straightforward implementation of adaptive-ICA for estimation of subject-specific components. Future work includes the development of iterative estimation schemes for data shared between sites. One of the advantages of decentralized analysis pipelines is that only intermediary statistics are passed between sites, and full patient records are never released across the network. These kinds of decentralized algorithms are “plausibly private” (Sarwate et al., 2014), due to the lack of directly identifiable records in the global data network. We believe work presented here provides a clear direction for future work in improving our understanding of brain disorders by leveraging data from different sites.

## 5 Conclusion

In this work, we demonstrated the feasibility of performing a multi-site large-sample ICA-based framework called Neuromark. The goal of Neuromark is to generalize and standardize the calculation of possible functional connectivity features that leverages the benefits of a data-driven approach and also provides comparability across multiple analyses. The analysis was performed on COINSTAC using functional MRI data collected in the UK and India. Additionally, we also demonstrated the decentralized statistical analysis of subject-specific network features for group level-inferences. We hope this work will be a useful stepping stone towards eventual application of such decentralized approaches for analyzing data from different sites and for further adoption in the clinical setting.

## 6 Conflict of interest

Dr. Banaschewski served in an advisory or consultancy role for Lundbeck, Medice, Neurim Pharmaceuticals, Oberberg GmbH, Shire. He received conference support or speaker’s fee by Lilly, Medice, Novartis and Shire. He has been involved in clinical trials conducted by Shire & Viforpharma. He received royalties from Hogrefe, Kohlhammer, CIP Medien, Oxford University Press. The present work is unrelated to the above grants and relationships. Dr. Barker has received honoraria from General Electric Healthcare for teaching on scanner programming courses. Dr. Poustka served in an advisory or consultancy role for Roche and Viforpharm and received speaker’s fee by Shire. She received royalties from Hogrefe, Kohlhammer and Schattauer. The present work is unrelated to the above grants and relationships. The other authors report no biomedical financial interests or potential conflicts of interest.

## 7 Data Availability Statement

Data sharing is not applicable to this article as no new data were created or analyzed in this study.

## 8 Information Sharing Statement

More specific details about accessing the datasets used in the study can be found at (Zhang et al., 2020) for cVEDA and (Schumann et al., 2010) for IMAGEN. The COINSTAC software can be accessed at https://github.com/trendscenter/coinstac.

## 9 Acknowledgments

This work was funded by the National Institutes of Health (grants: R01DA040487, R01MH121246, R01DA049238, and 1R01DA040487) and the National Science Foundation (grant: 2112455). This work received support from the following sources: the European Union-funded FP6 Integrated Project IMAGEN (Reinforcement-related behaviour in normal brain function and psychopathology) (LSHM-CT-2007-037286), the Horizon 2020 funded ERC Advanced Grant ‘STRATIFY’ (Brain network based stratification of reinforcement-related disorders) (695313), Human Brain Project (HBP SGA 2, 785907, and HBP SGA 3, 945539), the Medical Research Council Grant ’c-VEDA’ (Consortium on Vulnerability to Externalizing Disorders and Addictions) (MR/N000390/1), the National Institute of Health (NIH) (R01DA049238, A decentralized macro and micro gene-by-environment interaction analysis of substance use behavior and its brain biomarkers), the National Institute for Health Research (NIHR) Biomedical Research Centre at South London and Maudsley NHS Foundation Trust and King’s College London, the Bundesministeriumfür Bildung und Forschung (BMBF grants 01GS08152; 01EV0711; Forschungsnetz AERIAL 01EE1406A, 01EE1406B; Forschungsnetz IMAC-Mind 01GL1745B), the Deutsche Forschungsgemeinschaft (DFG grants SM 80/7-2, SFB 940, TRR 265, NE 1383/14-1), the Medical Research Foundation and Medical Research Council (grants MR/R00465X/1 - MRF-058-0004-RG-DESRI and MR/S020306/1 - MRF-058-0009-RG-DESR-C0759), the National Institutes of Health (NIH) funded ENIGMA (grants 5U54EB020403-05 and 1R56AG058854-01). Further support was provided by grants from: the ANR (ANR-12-SAMA-0004, AAPG2019 - GeBra), the Eranet Neuron (AF12-NEUR0008-01 - WM2NA; and ANR-18-NEUR00002-01 - ADORe), the Fondation de France (00081242), the Fondation pour la Recherche Médicale (DPA20140629802), the Mission Interministérielle de Lutte-contre-les-Drogues-et-les-Conduites-Addictives (MILDECA), the Assistance-Publique-Hôpitaux-de-Paris and INSERM (interface grant), Paris Sud University IDEX 2012, the Fondation de l’Avenir (grant AP-RM-17-013), the Fédération pour la Recherche sur le Cerveau; the National Institutes of Health, Science Foundation Ireland (16/ERCD/3797), U.S.A. (Axon, Testosterone and Mental Health during Adolescence; RO1 MH085772-01A1), and by NIH Consortium grant U54 EB020403, supported by a cross-NIH alliance that funds Big Data to Knowledge Centres of Excellence. ImagenPathways ”Understanding the Interplay between Cultural, Biological and Subjective Factors in Drug Use Pathways” is a collaborative project supported by the European Research Area Network on Illicit Drugs (ERANID). This paper is based on independent research commissioned and funded in England by the National Institute for Health Research (NIHR) Policy Research Programme (project ref. PR-ST-0416-10001). The views expressed in this article are those of the authors and not necessarily those of the national funding agencies or ERANID. Author ZZ received a fellowship from the Medical Research Foundation (MRF-058-0014-F-ZHAN-C0866).

## 10 Author Contributions

HG led the manuscript writing with support from KR and SB. BH contributed data and was instrumental in performing the analysis in COINSTAC. ZZ contributed data to the study as well as contributed to describing the data. EV and RK manage the COINSTAC project. SP, AD and JT are part of the COINSTAC team. VB and GS are the principal investigators of the CVEDA-IMAGEN consortium and have been instrumental in facilitating this multi-site study. VC is also a co-investigator and leads the COINSTAC team, formed the vision for this work, and helped interpret the results. All others who were not mentioned here are part of either the cVEDA or IMAGEN consortia.

